# Toxigenic and zoonotic *Corynebacterium ulcerans* emerges from widespread hedgehog disease

**DOI:** 10.1101/2020.07.17.208827

**Authors:** An Martel, Filip Boyen, Jörg Rau, Tobias Eisenberg, Andreas Sing, Anja Berger, Koen Chiers, Sarah Van Praet, Serge Verbanck, Muriel Vervaeke, Frank Pasmans

**Affiliations:** Ghent University, Merelbeke, Belgium; Chemical and Veterinary Analysis Agency Stuttgart, Fellbach, Germany; Hessian State Laboratory, Giessen, Germany; Bavarian Health and Food Safety Authority (LGL), Oberschleißheim, Germany; Agency for Nature and Forests, Belgium

## Abstract

Toxin-producing *Corynebacterium ulcerans*, causing diphtheria in humans, were isolated from 53 diseased hedgehogs during spring 2020 in Belgium. Isolates showed low levels of acquired antimicrobial resistance. Pronounced strain diversity suggests emergence from an endemic situation. These findings stress the need for raising public awareness and improved wildlife disease surveillance.

## Widespread occurrence of *Corynebacterium ulcerans* in hedgehogs with ulcerative skin disease

During May and June 2020, we obtained 60 isolates of *C. ulcerans* from 81 diseased or dead hedgehogs (*Erinaceus europaeus*) that were presented with ulcerative skin lesions by the public to four animal rescue centers across northern Belgium (*i*.*e*. Flanders) (Figure 1). The bacterium was isolated from ulcers or abscesses on the head or limbs of 53 hedgehogs, all of them being adult males. From six animals, more than one isolate was obtained from different lesions. Although *C. ulcerans* was most often isolated these lesions yielded abundant, poly-bacterial growth (Table 1). Systematic post mortem examinations of nine animals showed good body condition of the animals, with multiple cutaneous ulcers on the head and limbs. Histopathology from the skin of these animals revealed extensive ulcerative dermatitis with phlegmonous inflammation, sometimes extending in the subcutis and even underlying skeletal muscles. Intralesional bacteria as well as fragments of plant material were noticed. Four animals showed an interstitial pneumonia, one animal suffered from absceding hepatitis and in one animal, a fibrinosuppurative epicarditis with intralesional bacteria was noticed. All animals had extramedullar hematopoiesis in the liver and spleen. Despite presence of several other pathogens such as fly maggots (myiasis, n = 5), *Sarcoptes scabiei* (n=1), Herpesvirus (n=2), *Caparinia* spp. (n=1) and *Cresonema striatum* (n=2), no consistent evidence for other, primary disease causes was found. While evidence does not suffice to attribute a causal role for *C. ulcerans* in the lesions observed, its widespread and high-level occurrence in diseased male hedgehogs is worrying and constitutes a public health risk, given frequent exposure of humans to hedgehogs and because *C. ulcerans* is the predominant cause of human diphtheria in many European countries (*1*).

**Table 1:** Bacterial isolates obtained from 81 diseased hedgehogs across Flanders, Belgium. Numbers reflect the number of positive animals for the respective bacterium. Bacteria were identified using MALDI-TOF mass spectrometry. Biovar identification for the *C. diphtheriae* isolate was performed using the Api Coryne system (BioMerieux).

| Bacterial species obtained from lesions | Number of positive animals |
| --- | --- |
| *Corynebacterium ulcerans* | 53 |
| *Staphylococcus aureus* | 25 |
| *Enterococcus faecalis* | 23 |
| *Streptococcus canis* | 18 |
| *Streptococcus dysgalactiae* | 18 |
| *Proteus vulgaris/P. hauseri* | 17 |
| *Staphylococcus rostri* | 15 |
| *Proteus mirabilis* | 14 |
| *Streptococcus pyogenes* | 13 |
| *Staphylococcus xylosus* | 10 |
| *Staphylococcus microti* | 8 |
| *Staphylococcus sciuri* | 8 |
| *Vagococcus fluvialis* | 8 |
| *Enterococcus avium* | 7 |
| *Escherichia coli* | 7 |
| *Staphylococcus pettenkoferi* | 7 |
| *Morganella morganii* | 6 |
| *Staphylococcus fleurettii* | 6 |
| *Pasteurella multocida* | 4 |
| *Corynebacterium amycolatum* | 3 |
| *Bacteroides fragilis* | 3 |
| *Staphylococcus simulans* | 3 |
| *Trueperella pyogenes* | 3 |
| *Vagococcus lutrae* | 2 |
| *Arcanobacterium haemolyticum* | 1 |
| *Bacillus* sp. | 1 |
| *Bacteroides pyogenes* | 1 |
| *Corynebacterium confusum* | 1 |
| *Corynebacterium diphtheriae mitis* | 1 |
| *Enterococcus hirae* | 1 |
| *Enterobacter hormaechei* | 1 |
| *Gemella haemolysans* | 1 |
| *Lactococcus garvieae* | 1 |
| *Streptococcus gallinaceus* | 1 |
| *Streptococcus thoraltensis* | 1 |

**Figure 1.**
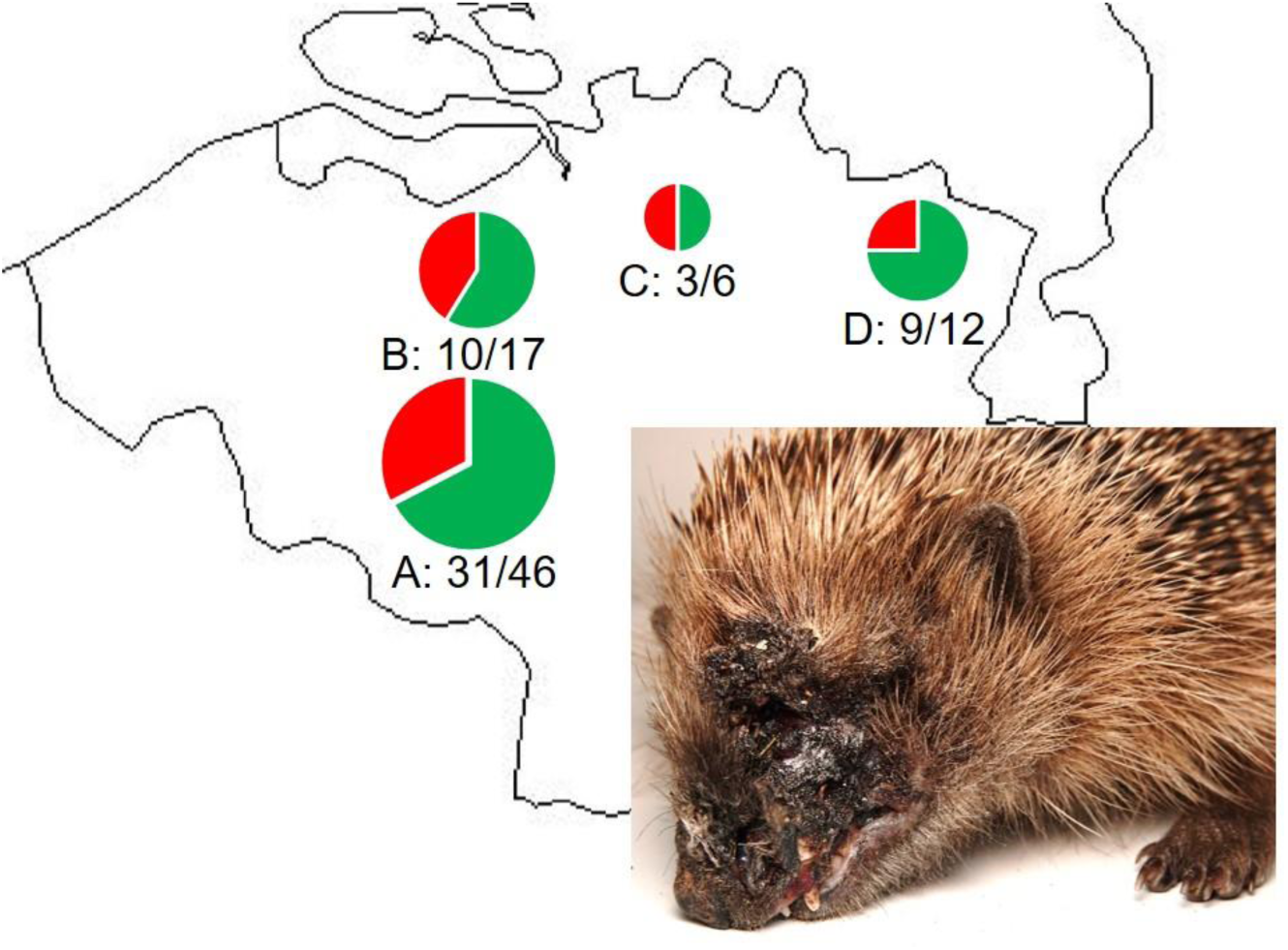
Eighty-one hedgehogs with lesions on the head or limbs from four regions in Flanders were investigated for the presence of *C. ulcerans* (green: proportion of positive animals; red: proportion of negative animals). Regions A: Geraardsbergen, B: Merelbeke, C: Herenthout, D: Oudsbergen. Small picture indicates a representative clinical state with necrotizing facial dermatitis in one male hedgehog from Merelbeke.

## *Corynebacterium ulcerans* isolates from hedgehogs belong to several clusters

Fifty-six isolates of *C. ulcerans*, identified on species level using MALDI-TOF MS (*2*) and sequencing of the *rpoB* gene [3], were typed by analysis of infrared spectra (*4*). The isolates grouped with *C. ulcerans* strains from humans and other animals (hedgehogs and red foxes from Germany) (*5, 6*) and clustered in three sublineages (Figure 2). Results argue against emergence and spread of a single *C. ulcerans* clone in the hedgehog population in Flanders. Instead, the high diversity of the isolates suggests *C. ulcerans* endemism in the hedgehog population.

**Figure 2.**
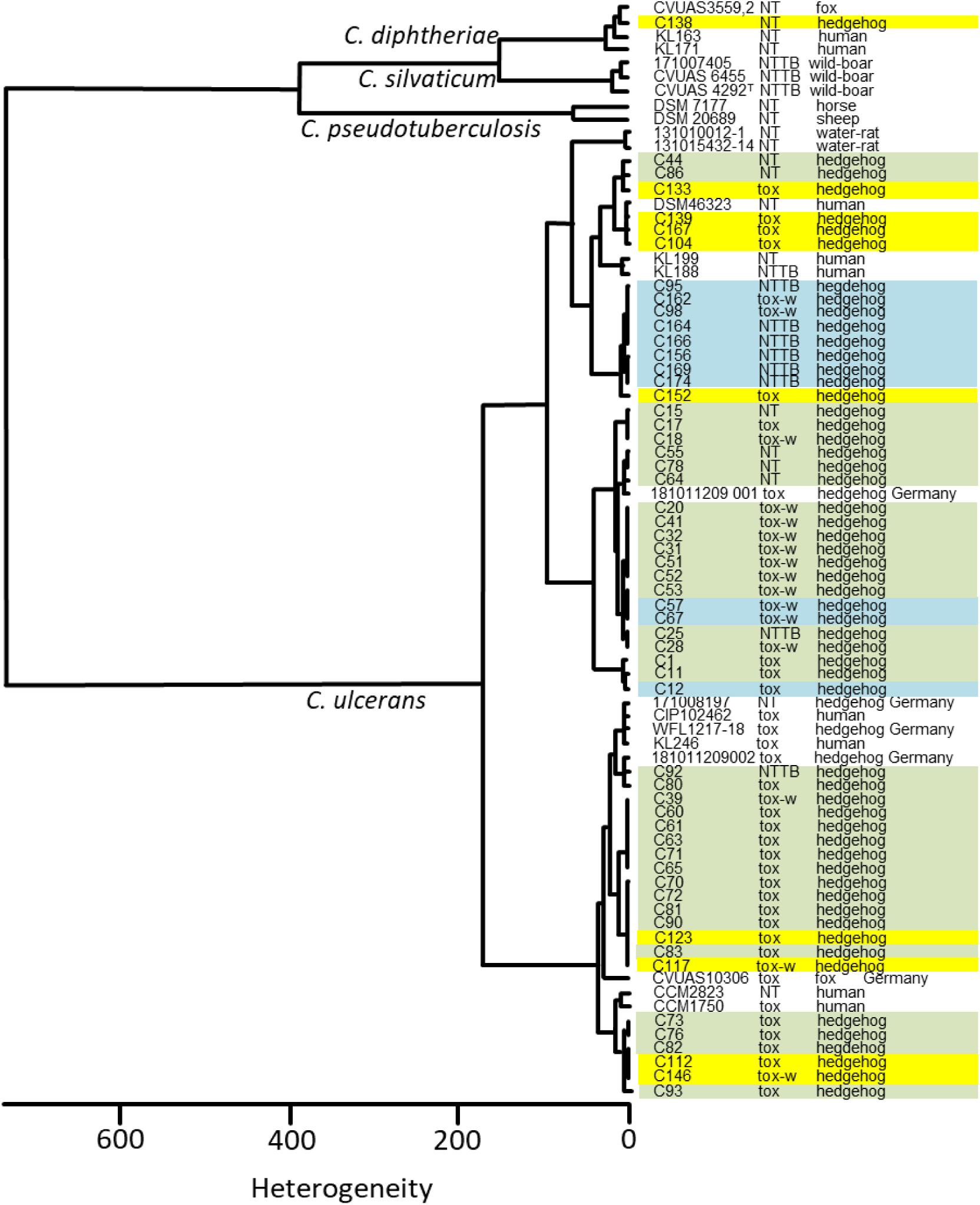
Dendrogram of the FT-IR spectra of C*orynebacterium* sp. strains obtained from hedgehogs in comparison with spectra from several *C. ulcerans* isolates, including isolates from other free roaming animals and humans [cf. 8]. Region: Geraardsbergen, green; Merelbeke, blue; Oudsbergen, yellow. tox: toxigenic; tox-w: toxigenic (weak), NTTB: non toxic but tox bearing; NT: no *tox*-gen. Further details of the isolates are shown on MALDI-UP (*7*).

## Limited indications of acquired antimicrobial resistance in *Corynebacterium ulcerans* isolates from hedgehogs

Minimal inhibitory concentration (MIC) data of all *C. ulcerans* isolates are presented in Table 2. Acquired resistance against enrofloxacin and spiramycin was noticed in 4 and 1 isolates respectively.

**Table 2.** Distribution of MIC (µg/ml) values in the 60 isolates of *Corynebacterium ulcerans* isolates from this study as obtained by epsilometer (ETEST, bioMérieux, Marcy l’Etoile, France) on Mueller–Hinton agar plates with 5 % horse blood and 20 mg l^−1^ β-NAD (bioMérieux) according to both European Committee on Antimicrobial Susceptibility Testing (EUCAST) and Clinical and Laboratory Standards Institute (CLSI) guidelines (*8, 9*).

|  | <0.016 | 0.016 | 0.032 | 0.064 | 0.12 | 0.25 | 0.5 | 1 | 2 | 4 | 8 | 16 |
| --- | --- | --- | --- | --- | --- | --- | --- | --- | --- | --- | --- | --- |
| Amoxicillin | 1 | 10 | 6 | 8 | 25 | 10 |  |  |  |  |  |  |
| Amoxicillin-clavulanate | 2 | 2 | 2 | 12 | 18 | 22 | 2 |  |  |  |  |  |
| Clindamycin |  |  |  |  |  |  |  | 34 | 26 |  |  |  |
| Doxycyclin |  |  |  |  | 5 | 27 | 13 | 13 | 2 |  |  |  |
| Enrofloxacin |  | 4 | 33 | 19 |  |  |  | 4 |  |  |  |  |
| Erythromycin |  | 1 | 50 | 9 |  |  |  |  |  |  |  |  |
| Penicillin | 1 | 1 | 4 | 6 | 17 | 20 | 11 |  |  |  |  |  |
| Spiramycin |  |  |  | 43 | 17 |  |  |  |  |  |  |  |
| Tetracyclin |  |  |  |  |  | 11 | 29 | 19 | 1 |  |  |  |
| Tilmicosin |  |  |  |  |  |  | 1 | 29 | 29 | 1 |  |  |
| Trimethoprim sulfamethoxazole |  |  | 9 | 43 | 8 |  |  |  |  |  |  |  |

## Most *Corynebacterium ulcerans* isolates from hedgehogs produce toxins

Presence and expression of toxicity were evaluated by detection of the diphtheria toxin gene (*toxE*) using a duplex PCR (*3*) and the Elek test (*10*). Results showed the presence of the *toxE* gene in 50/56 isolates and a positive or weak positive Elek test in 26/56 and 16/56 isolates, respectively. One animal carried a *toxE* gen positive (isolated from a head lesion) and *toxE* gene negative (isolated from a foot lesion) *C. ulcerans* isolate. Since the presence of the *toxE* gene and the toxin production are associated with pathogenicity in humans, these results suggest a zoonotic risk of most hedgehog derived *C. ulcerans* isolates.

## Public and animal health risk of ulcerative hedgehog disease

Hedgehogs are abundant mammals in Europe, frequently observed in both nature reserves and urbanized areas. Due to their defensive behavior, sick animals are easily presented to animal rescue centers by the public, as testified by the large number of animals we examined in a short time frame in this study. The very nature of their spiny defense promotes breaching and inoculating the human epidermis with bacteria during handling. As demonstrated in Table 1, several potentially zoonotic bacterial species, including *C. diphtheriae* and *Streptococcus pyogenes* are associated with ulcerative lesions in diseased hedgehogs. However, it is the widespread occurrence of toxigenic *C. ulcerans* in the majority of the diseased hedgehogs across Flanders that should prompt authorities to alert all stakeholders, including the public and the animal rescue centers, to take precautionary measures when handling hedgehogs. While vaccination against *C. diphtheriae* protects against *C. ulcerans* disease, exposure to *C. ulcerans* of susceptible people may result in severe disease (*11*). Recommendations should include wearing protective gloves and cleaning and disinfecting hands and fomites after contact with a hedgehog and vaccination of people who are frequently exposed to hedgehogs. Treatment of infections with pyogenic coryneform bacteria in animals is challenging and the four rescue centers involved in this study reported poor treatment success. The results of the antimicrobial susceptibility testing suggest that this is not due to acquired antimicrobial resistance, but probably rather to insufficiently high antimicrobial concentrations reaching the *C. ulcerans* bacteria inside pus. Debridement of the lesions should, therefore, be included in any treatment. Euthanasia should be considered for severe cases.

Emergence of *C. ulcerans* in hedgehogs is in line with a growing number of reports of *C. ulcerans* infections in wildlife across Europe and warrants attention across the continent (*5, 6, 12, 13, 14*). Reports dating back from the 1950s and the presence of several distantly related clusters of *C. ulcerans* in this study argue against a recent introduction of this pathogen in European wildlife populations and favors the hypothesis that the observed and previously unreported, high numbers of diseased hedgehogs result from pathogen emergence from an endemic state. The finding that only male hedgehogs were presented with this disease, and lesions are mostly found on body parts not covered with spines, suggests the *C. ulcerans* infections may be opportunistic infections of wounds, arising from male specific behavior during the mating season. Bite wounds are well known to be prone to infection with opportunistic pathogens that are part of the oral microbiota (*15*). The strain typing indeed suggests that hedgehogs constitute a significant reservoir of highly diverse *C. ulcerans* isolates. Active surveillance should elucidate the magnitude of this reservoir in healthy hedgehogs and other wildlife. Further studies should unravel the mechanisms underpinning the observed emergence of this zoonotic wildlife disease from its endemic state.

## Acknowledgements

This work was funded by the Agency for Nature and forests of the Flemish government, the MALDI-TOF MS was financed by the Research Foundation Flanders (FWO-Vlaanderen) as a Hercules project (G0H2516N, AUGE/15/05), the BOF research as a ZAP mandate to A. Martel and the Bavarian State Ministry of Health and Care as well as by the German Federal Ministry of Health via the Robert Koch-Institute (09-47, FKZ 1369-359). We thank the rescue centers VOC Wilde Dieren (Nancy Van Liefferinge and Filip Berlengee), VOC Merelbeke (Nick De Meulemeester), Natuurhulpcentrum (Frederick Thoelen) and VOC Neteland (via Tom Verbeek) for the participation to this study.

